# Adaptive energetic tuning of nucleolar phase separation regulates rRNA transport

**DOI:** 10.64898/2026.08.24.746772

**Authors:** Evdokiia Potolitsyna, Elmira Khabusheva, Yiling Fang, Erik Bergstrom, Hazel A. Mangan, S. Thomas Hennigan, Zion Ibarra, Christina Dollinger, Nayara Alcantara-Contessoto, Margaret A. Goodell, Lucia Strader, Brian McStay, Joshua A. Riback

## Abstract

Eukaryotic cells adjust biochemical pathways in response to environmental stressors, stalling energetically costly processes such as protein synthesis and ribosome biogenesis. Ribosome biogenesis occurs within the multiphase nucleolus that contains three layers roughly corresponding to ribosomal RNA (rRNA) transcription, processing, and assembly. While nucleolar perturbations induced by drug treatments and optogenetic nucleolar gelation alter nucleolar stability and hinder ribosome biogenesis, it remains unclear whether nucleolar properties are actively tuned in response to metabolic and growth cues. Here, we show that nucleolar phase separation is adjusted to physiological needs. Live-cell imaging of endogenously tagged NPM1 reveals a tight relationship between NPM1 partitioning and nutrient availability. In nutrient-deprived conditions, NPM1 levels in the nucleolus increase, indicative of stabilized phase separation and a more gel-like state. Indeed, we show that rRNA diffusion decreases and the nucleolar meshwork contracts. Furthermore, this change corresponds to decreased rRNA processing, suggesting that reduced transport properties prevent the release of immature ribosome subunits from the nucleolus. Mechanistically, we show that these changes are driven by ATP levels, which broadly affect the energetically expensive process of ribosome biogenesis. Upon restoration of ATP levels by nutrient reintroduction, nucleolar composition returns to normal within minutes, suggesting active regulation. Taken together, our findings reveal that the biophysical properties of the nucleolus are not fixed, but are actively remodeled by cellular energy levels, linking phase separation dynamics to metabolic control of ribosome biogenesis.

## Introduction

Eukaryotic cells regulate biochemical pathways in response to various environmental stressors, often stalling key processes such as RNA transcription and protein synthesis^1,2,8^. Ribosome biogenesis is energetically expensive, requiring coordinated transcription by RNA polymerases I, II, and III, as well as the synchronized production of 80 ribosomal proteins^9,10^. Indeed, ribosomal content is tightly coupled with cell growth, and under stress or energy limitation, ribosome biogenesis must therefore be rapidly adjusted to cellular demand^11^.

Ribosomes originate from the nucleolus, where ribosomal RNA (rRNA) is synthesized, and pre-ribosomal subunits are assembled^12^. The nucleolus consists of three subcompartments: the fibrillar center (FC), the dense fibrillar component (DFC), and the granular component (GC), which spatially coordinate the sequential steps of rRNA transcription, processing, and ribosomal subunit assembly^3^. Multiple imaging studies cataloged how nucleolar morphology responds to various environmental stressors and drug perturbations, a phenomenon broadly termed nucleolar stress, which is often accompanied by translocation of the major nucleolar protein Nucleophosmin 1 (NPM1) from the nucleolus to the nucleoplasm^4,6,7,13,14^. However, morphological changes alone provide little insight into the precise impact of the nucleolus on the regulation of ribosome biogenesis.

The emergence of phase separation as a framework for nucleolar formation has recast the nucleolus as a multiphase, viscoelastic material in which organization, composition, material state, and ribosome biogenesis are inherently coupled^5,15–18^. For instance, inhibition of rRNA transcription with Actinomycin D progressively weakens nucleolar phase separation as ribosomal intermediates continue to mature and leave the nucleolus in the absence of newly synthesized rRNA^19,20^. This indicates that as pre-ribosomal particles gradually fold, they lose valency for nucleolar interactions, ultimately leading to their thermodynamic exclusion from the nucleolus^19–21^. Consistent with this, early pre-ribosomal particles form a tighter, more stable meshwork and move more slowly through the nucleolus^15,22^. In contrast, optogenetically induced nucleolar gelation impairs rRNA processing^5^ and targeting an engineered killswitch peptide to the nucleolus arrests nucleolar dynamics, disrupting nucleolar function^23^. These studies, along with recent work mapping rRNA processing and flux across nucleolar phases^24,25^, suggest a broader condensate structure-function relationship whereby the biophysical properties of nucleolar phase separation directly impact the speed and accuracy of ribosome biogenesis. However, if these properties are dynamically tuned in response to physiological stimuli such as energy availability remains largely unclear.

Here, we show that nucleolar properties are dynamically tuned in living cells in response to nutrient availability. Using live-cell imaging of endogenously tagged nucleolar proteins, we find that serum and nutrient deprivation drive opposite changes in nucleolar NPM1 concentration and partitioning. Given NPM1’s role as an abundant GC scaffold, these changes indicate that GC phase behavior is modulated by nutrient availability. We characterize the nutrient-deprivation response using three orthogonal methods that all show nutrient deprivation causes a less dynamic, more viscous nucleolar state in which rRNA maturation and transport through the nucleolus are slowed. We define this response as nucleolar “hibernation”. Nucleolar hibernation is rapidly reversible upon refeeding and is coupled to cellular ATP levels, indicating active metabolic control. Comparable stabilization of nucleolar phase separation in mouse hematopoietic stem cells and *Nicotiana benthamiana* leaf epidermal cells suggests that energy-dependent tuning of nucleolar organization represents a conserved adaptive response that coordinates ribosome biogenesis with cellular metabolic state. More broadly, our findings support a role for phase separation not only in compartmentalizing biochemical reactions, but also in tuning their output to physiological demand.

## Results

### NPM1 enrichment in the GC is a fine-tuned response to nutritional stresses

To investigate whether nucleolar composition can be fine-tuned, we focused on nutrient availability as a model perturbation. Nutrient availability is a physiologically relevant parameter of cellular homeostasis and is straightforward to titrate experimentally. Depletion of FBS targets primarily mitogenic signaling and is often used to synchronize cell cultures in the G0/G1 stage, whereas graded depletion of all major nutrients induces broader metabolic adaptation^26,27^. We subjected U2OS cells to different media conditions and performed immunofluorescence staining with antibodies against NPM1, observing depletion of NPM1 from the nucleoli in both the serum-deprived (SD) and nutrient-deprived (ND) conditions (**Fig. 1A**). Indeed, multiple previous studies using immunofluorescence show depletion of NPM1 in the nucleoli upon diverse stressors, including various starvation conditions, commonly referred to as translocation of NPM1 into the nucleoplasm^13,28,29^.

**Figure 1.**
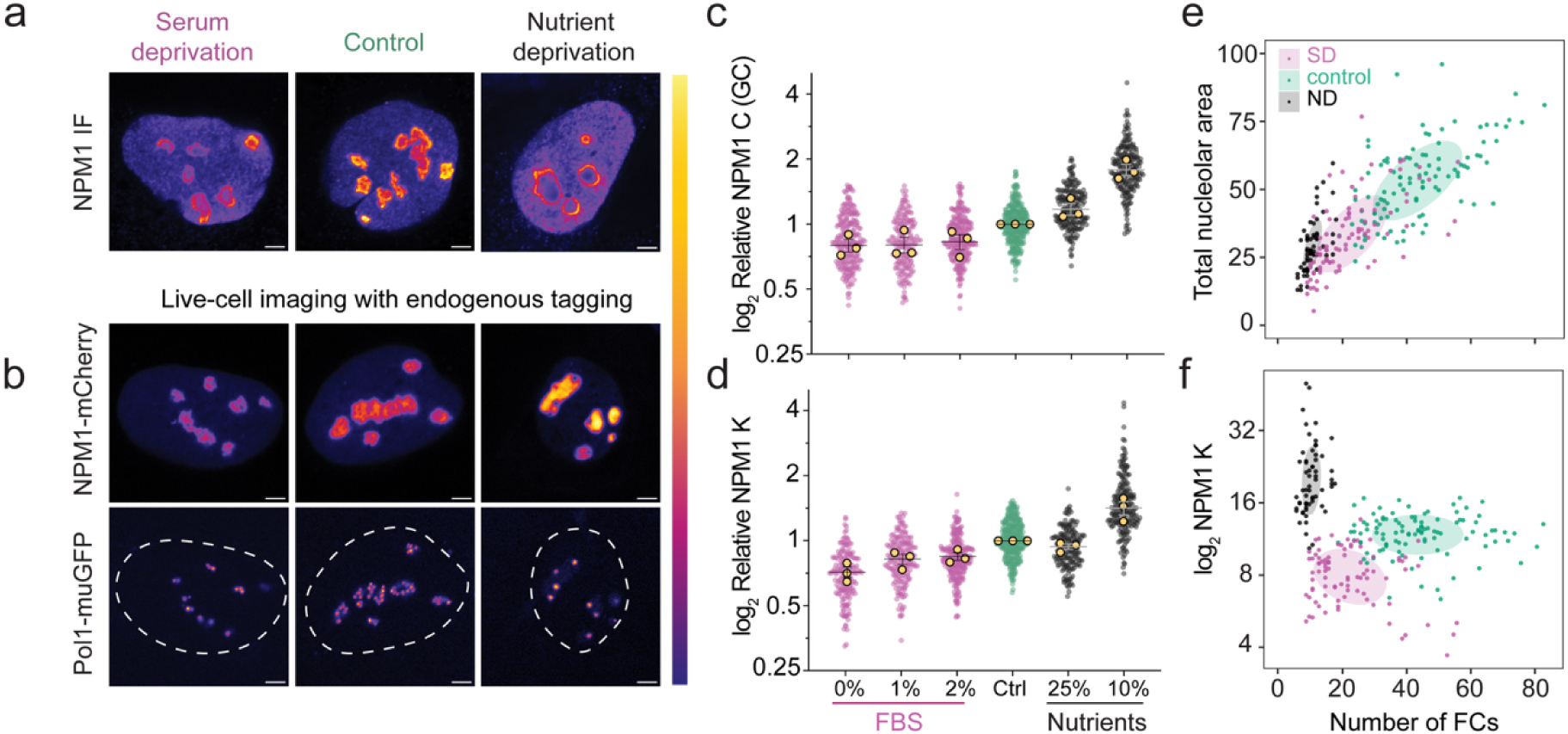
Nutrient deprivation and serum withdrawal induce distinct nucleolar responses. (**A**) Immunofluorescence images of U2OS cells stained with antibodies against NPM1. (**B**) Live-cell imaging of U2OS cells with endogenously edited NPM1-mCherry and Pol1-muGFP. (**C**) Relative levels of NPM1 in the GC in U2OS cells from (B), normalized to the control condition, n = 3 experiments with N > 35 cells per condition per replicate (gold circles); data is presented as mean ±S.E.M. (**D**) Relative partitioning of NPM1, similar to (C). (**E**) Scatter plot of total nucleolar area and (**F**) NPM1 partitioning versus the number of FCs per cell; N > 50 cells per condition; shaded regions correspond to 68% confidence region. Images are shown using fire LUTs, with nucleoplasmic levels fixed across individual panels; scale bars 3 µm.

To understand the underlying changes in interactions that lead to NPM1 depletion from the nucleolus, we generated a CRISPR knock-in line expressing NPM1-mCherry, enabling us to study NPM1 behavior in living cells (**Fig. S1A**). In contrast to IF staining, nucleoli of these cells exhibited visually distinct responses, with an increase in NPM1 in the nucleoli of cells deprived of 90% of nutrients (ND) (**Fig. 1B**), including in edited cells co-stained with an antibody (**Fig. S1B**). We then turned to partitioning measurements to quantitatively assess the thermodynamic strength of NPM1 transfer from the nucleoplasm to the nucleolus under titrated nutrient-deprivation conditions (**Fig. 1C, D**). Removal of FBS resulted in a gradual decrease of NPM1 concentration in the nucleolus, an increase of NPM1 in the nucleoplasm, and a drop in NPM1 partitioning to 0.7±0.1 fold at 0% FBS (**Fig. 1D**). Surprisingly, titrated removal of all nutrients (ND) led to a gradual increase of NPM1 concentration in the nucleolus, reaching 1.8±0.2 fold relative to the control condition. At the same time, NPM1 partitioning increased from 14±1 to 20±2 (1.5±0.2 fold) at 10% nutrients (**Fig. 1D**). Such increases under ND indicate a substantial remodeling of the interactions stabilizing the GC.

GC-remodeling could involve either increased or decreased nucleolar activity (rRNA transcription). To distinguish these, we measured the total number of FCs per cell, as marked by endogenously labeled Pol1-muGFP (**Fig. 1B**, lower; Fig. S1A), a longstanding measure of nucleolar activity^30^. Both serum deprivation and ND decrease the total number of FCs, indicative of lower rRNA production (**Fig. 1E**). Additionally, SD is accompanied by reduced NPM1 partitioning and nucleolar area, suggesting weakened preferential interactions between NPM1 and rRNA and weaker phase separation within the nucleolus (**Fig. 1D, F**), in line with our previous observations under acute inhibition of rRNA production by Actinomycin D^19^. In contrast, ND causes an increase in NPM1 partitioning that clearly distinguishes it from SD (**Fig. 1F**). Because serum and nutrient deprivation can alter cell-cycle distribution, we tested whether these distinct nucleolar responses could be explained by cell-cycle responses, finding that the enrichment of SD cells in G1 (6.3%) and ND cells in G2 (6%) compared to control was insufficient to account for their effects on NPM1 partitioning, nucleolar area, or FC number (**Fig. S1C**). These findings suggest that ND uncovers a previously unrecognized axis of nucleolar regulation, in which GC organization and thermodynamics can be modulated independently of transcriptional activity.

### Nutrient deprivation triggers nucleolar “hibernation” response, characterized by slower processing of rRNA

The increased thermodynamic stabilization of NPM1 in ND does not resemble previously described nucleolar phenotypes that generally feature NPM1 depletion from the GC and nucleolar dissolution^4,6,7,13,14^, prompting us to investigate this distinct nucleolar state further. Nucleolar properties are strongly defined by the RNA component, which contributes to nucleolar viscoelasticity^15,17^. Additionally, the extent of NPM1 enrichment into the GC is suggested to be dictated by rRNA folding during ribosome biogenesis^19–21^. Thus, we evaluated the status of rRNA transcription and processing in cells under control and nutrient-deprived conditions with qPCR (**Fig. 2A-C**). We designed primers targeting various regions of the rRNA transcript, allowing us to distinguish nascent rRNA, total nucleolar intermediates, pre-GC intermediates, and early GC intermediates, respectively (**Fig 2A-B**, Methods). In agreement with the decrease in the number of FCs per cell (**Fig. 1E, F**), transcription of nascent 47S rRNA is reduced by ∼70-90% in ND (**Fig. 2C**). However, the total amount of rRNA intermediates does not significantly change, suggesting their retention and a slower rate of processing. Specifically, pre-GC rRNA intermediates decreased by 30%, whereas early GC intermediates showed no significant change (**Fig. 2C**). When taking into account the halving of nucleolar size during ND (**Fig. 1E**), this implies a doubling of the concentration of early GC intermediates. Overall, these results show that ribosome biogenesis is heavily remodeled during ND and that despite the decreased transcription of rRNA, rRNA intermediates accumulate and concentrate within the GC. The combination of stalled ribosome biogenesis and altered nucleolar thermodynamics defines a nucleolar response distinct from other low-activity responses, such as SD, and rRNA transcription inhibitors. As rRNA intermediates and NPM1 seem sequestered, possibly to maintain nucleolar integrity, we will refer to this response as *nucleolar hibernation* and further investigate the biochemical and biophysical features that regulate the transition into and maintenance of nucleolar hibernation.

**Figure 2.**
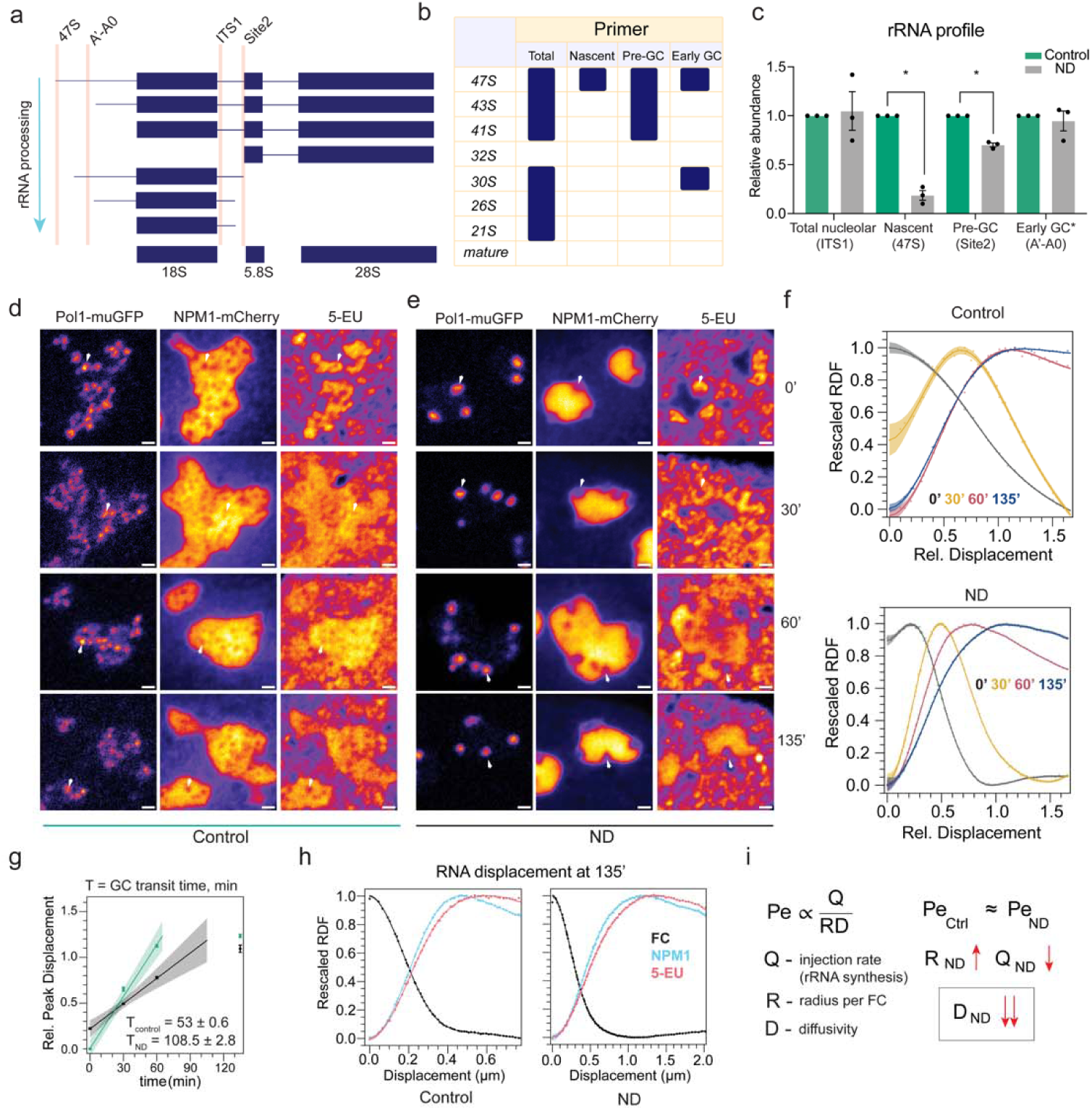
Nutrient deprivation slows rRNA maturation and progression through the granular component. (**A**) Schematic of qPCR primer placement on rRNA precursors. (**B**) Table summary of primer-targeted intermediates. (**C**) Relative levels of rRNA intermediates in control and nutrient-deprived cells, normalized to control (mean fold change ±S.E.M.; paired t test, n = 3; *p<0.05). (**D-E**) 5-EU pulse-chase time course in (D) control and (E) ND cells. (**F**) Corresponding radial distribution functions (RDFs) for 5-EU signal plotted relative to the NPM1 peak; N ≥ 10 cells per time point. (**G**) RNA peak displacement over time and estimated GC transit time based on (F). (**H**) RDF plot for a 135-minute chase with the addition of FC and GC signals, used to estimate the Peclet number. (**I**) Peclet number scaling with rRNA injection rate (Q), GC radius (R), and diffusivity (D), suggesting reduced diffusivity under ND conditions. Images are shown using fire LUTs; scale bars: 1 µm.

### Hibernating nucleoli exhibit restricted rRNA transport dynamics

As rRNA intermediates mature, they progressively move outward from the site of transcription (FC) ^15,24,25^. Notably, we previously found a role for advection in rRNA transport within the GC that maintains a peaked movement^15^. Thus, we asked how nucleolar hibernation impacts rRNA transport and advective flow. To measure rRNA transport through the nucleolus, we performed 5-EU labeling of nascent RNA and quantified the radial distribution function (RDF) in the nucleolus from the center of the FCs, where rRNA transcription occurs (**Fig. 2D,E; Fig. S2A**). Incorporation of 5-EU over 30 minutes reveals a low but detectable level of rRNA transcription in cells exposed to ND (**Fig. 2E**), in agreement with qPCR data showing low levels of nascent rRNA (**Fig. 2A-C**, 47S primer). After 30 minutes of chase in control cells, the labeled RNA is organized in a tight migrating front in agreement with previous data (**Fig. 2F, Fig. S2B**)^15,24,25^. By 60 minutes chase in control cells, the labeled RNA had reached its maximal radial extent, and the distributions at 60 and 135 minutes largely overlapped with the NPM1 peak, indicating steady-state equilibration within the GC (**Fig. 2F, Fig. S2B**). In contrast, nutrient-deprived cells exhibited a clear delay in this progression, with continued displacement of the labeled RNA between the 60- and 135-min time points (**Fig. 2E, F**). We measured peak displacement at each chase time point and defined the GC transit time as the average time required for newly synthesized rRNA to transit through the GC (**Fig. 2G**, Methods). The GC transit time in control cells is 53±1 min, consistent with that reported in HEK cells^15^. In contrast, rRNA movement in the hibernated nucleolus had a GC transit time of 109±3 min, a near doubling.

The doubling of GC transit time during ND is particularly notable because slower passage through the GC should allow diffusion to broaden the rRNA front. Yet the rRNA RDF profiles in ND remain peaked throughout the time course, indicating that directed transport is preserved despite the greater transit time. To better quantify the balance between advection (directed flow) and diffusion, we measured the apparent Peclet number (Pe), a common measure of this balance, at the latest chase time point (**Fig. 2H**), as done previously^15^. We found a slight decrease in the Pe from 2.6±0.1 in control cells to 1.2±0.1, indicating that advection contributes slightly more than diffusion in both despite substantial architectural remodeling (**Fig. 2H**). Combined with the decrease in rRNA transcription-driven influx (Q, **Fig 2C**) and increase in transport distance (R; GC radius per FC, 0.47±0.01 µm vs. 1.22±0.04 µm), the similar Peclet number implies a reduction in the diffusivity of rRNA, indicating an increase in the viscosity of the GC (**Fig. 2I**). Together, these findings suggest that the hibernating nucleolus maintains directed rRNA transport despite extensive architectural remodeling, through coordinated changes in RNA flux and effective diffusivity.

We sought to confirm this conclusion using orthogonal methods. Notably, 5-EU pulse-chase experiments report on the average transport of nascent RNA from all active rDNA-containing chromosomes and require cell fixation. To achieve live-cell spatiotemporal mapping of rRNA transport dynamics, we next employed a previously described MS2-based live-cell reporter system based on an engineered RPE1 cell line containing an additional copy of chromosome 15, in which each 28S gene harbors a single MS2 hairpin repeat^31^ (**Fig. 3A**). We introduced stable expression of MS2-binding protein fused to Halo (MCP-Halo) in these cells and endogenously tagged NPM1 at its C-terminus with mStayGold, revealing that the MS2-containing chromosome occupies a distinct territory within the nucleolus (**Fig. 3A-B**), as previously described^31^. We further confirmed that MS2-containing rRNA progresses to the mature state and that NPM1 characteristics are unaffected in cells with the MS2 chromosome when compared to parental RPE1 (**Fig. S3A-B**).

**Figure 3.**
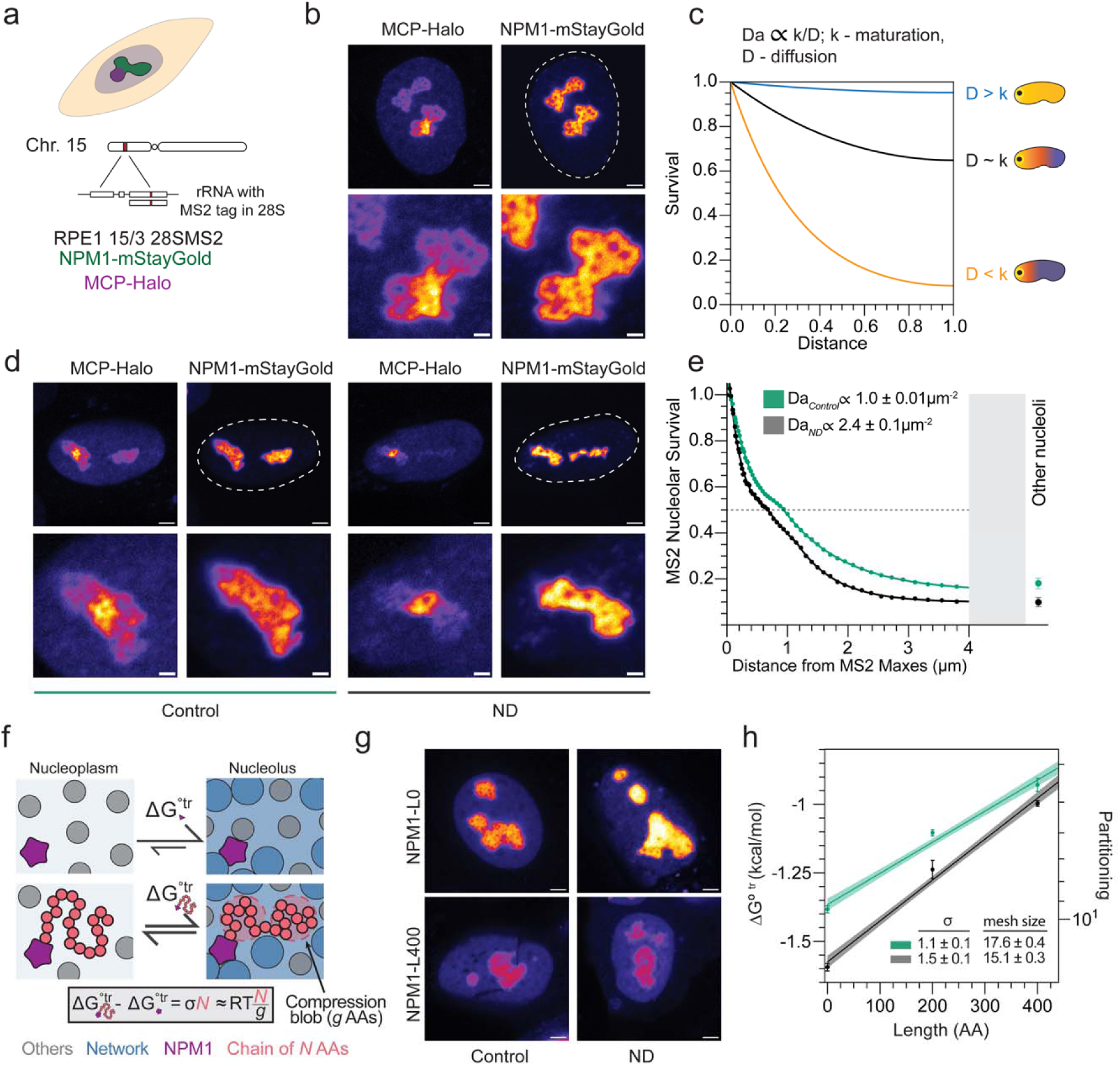
Hibernating nucleoli exhibit restricted rRNA mobility and a denser microenvironment. (**A**) Schematic of edited RPE1 HTERT cell line carrying an extra copy of chromosome 15 containing MS2-tagged rDNA repeats. (**B**) Live-cell imaging of the RPE1 cells with MCP-Halo labeling the rRNA transcribed from the MS2-containing chromosome; inset for the nucleolus at the bottom. (**C**) Damköhler number (Da) relationship and analytical model of expected rRNA signal decay under different relationships between reaction and transport rates. (**D**) RPE1 cells from (B) subjected to nutrient deprivation; inset for the nucleolus at the bottom. (**E**) MCP-Halo labeled rRNA signal decay within the nucleolus as a function of distance from each position of maximum MS2 intensity, averaged and fit to multi-exponential showing 50% peak decay and final decay length scale as proportionality to Da; maximal enrichment of labeled rRNA in other nucleoli not containing the MS2 chromosome (right); parameter standard error; N ≥ 25 cells per condition. (**F**) Schematic of the Local Size Exclusion method used to estimate the apparent mesh size surrounding NPM1 in the nucleolus. (G) Overexpression of NPM1, with or without the extra chain, in control and nutrient-deprived cells. (H) Partitioning of NPM1 constructs with or without additional disordered chains in control and nutrient-deprived cells; fit-propagated uncertainty; N ≥ 36 cells per data point; σ-value, kcal/mol/AA; mesh size, nm. All images are displayed in fire LUT, and scale bars are 3 µm or 1 µm for enlarged panels (B, D).

Outside of the chromosomal territory, where local advective flows from multiple FCs approximately cancel, the spatial distribution of MS2-labeled rRNA reflects both diffusive movement of ribosomal intermediates within the nucleolus and the rate-limiting maturation step that precedes rapid nucleolar exit. This reaction-transport balance is captured theoretically by the Damköhler number (Da), a dimensionless ratio of the effective reaction rate to the transport rate that governs signal survival^32^ (**Fig. 3C**). To analyze the MS2 signal, we quantified its radial decay within the nucleolus by measuring signal survival as a function of distance from each position of maximum intensity, averaged across nucleoli and fit (**Fig. 3D-E**, methods). The signal was more constrained during ND, with 50% MS2 survival occurring at 0.67±0.02 μm compared with 0.94±0.01 μm in control. Furthermore, at large distances, where the nucleolar multiphase structure is averaged out, the decay profile could be well described by a single exponential whose slope is proportional to the Da (**Fig. 3B**). Fitting yielded a 2.4±0.1 μm^-2^ during ND compared with 1.00±0.01 μm^-2^ in control conditions. The MS2 survival stalled to 0.146±0.001 and 0.099±0.001 at long distances for normal and hibernating nucleoli. These values were consistent with those observed in other nucleoli within the cell (**Fig. 3B**). While MCP is an RNA-binding protein, it does not preferentially localize in the nucleoli of RPE1 cells that do not contain the MS2 chromosome, confirming that the MCP signal detected in other nucleoli represents MS2-containing rRNA intermediates that diffused from the original nucleolus (**Fig. S3C**). Altogether, these four measures extracted from the MS2 signal establish that the Damköhler number increases during ND despite slower maturation of rRNA intermediates within the nucleolus, indicating, in agreement with the 5’-EU pulse-chase data, a significant decrease in diffusivity and thus increase in nucleolar viscosity.

### Nucleolar hibernation stabilizes a denser nucleolar microenvironment

To investigate whether restricted rRNA transport in hibernating nucleoli is directly associated with altered organization of the dynamic interaction network between nucleolar scaffold proteins and ribosomal intermediates, we employed Local Size Exclusion (LSE), a novel method that measures the local meshwork around individual proteins within a condensate^22^ (**Fig. 3G**). This method measures how the partitioning energy changes with the addition of a flexible amino acid chain, yielding a slope per amino acid (σ-value) that reports on the local confinement experienced by the chain within a condensate. Overexpressing NPM1 and subjecting cells to ND, we observed an increase in the partitioning of NPM1 without an extra chain, consistent with our data for endogenously tagged NPM1 (**Fig. 3G, H**; see also **Fig. 1B, D**). However, the sensitivity of NPM1 partitioning to the addition of an inert chain was greater during ND than control, corresponding to an increase in σ-value from 1.1 ± 0.1 to 1.5 ± 0.1 cal mol^-1^ per residue (**Fig. 3H; Fig. S3D**). This equates to a ∼15% decrease in the average effective local mesh size around NPM1, from 17.6±0.4 nm to 15.1±0.3 nm. Together with the 5-EU pulse-chase and MS2 results, these findings indicate that nucleolar hibernation produces a denser and more viscous nucleolus that slows RNA transport seemingly adapting the spatial release of ribosomal intermediates with their maturation kinetics.

### Nucleolar hibernation is fully reversible within minutes of nutrient re-addition

Transition to a hibernated nucleolar state raises the question of whether this represents a terminal, dysfunctional phenotype, as is often implicated for condensate gelation (i.e., irreversible gelation^33,34^), or instead is a form of cellular adaptation. Thus, we sought to examine whether nucleolar organization and dynamics can be restored upon nutrient re-addition. Reintroduction of nutrients in ND cells, resulted in nucleoli visually responding within minutes (**Fig. 4A-B; Fig. S4A**). Following individual cells on shorter timescales revealed that nucleolar NPM1 levels recover within minutes, with a half-time of 3.8 min (**Fig. 4B-C; Fig. S4B**), suggesting recovery of normal nucleolar function. These data show that nucleolar hibernation is rapidly reversible, supporting it as an adaptive state, not a dysfunctional one, that poises the nucleolus for rapid restart.

**Figure 4.**
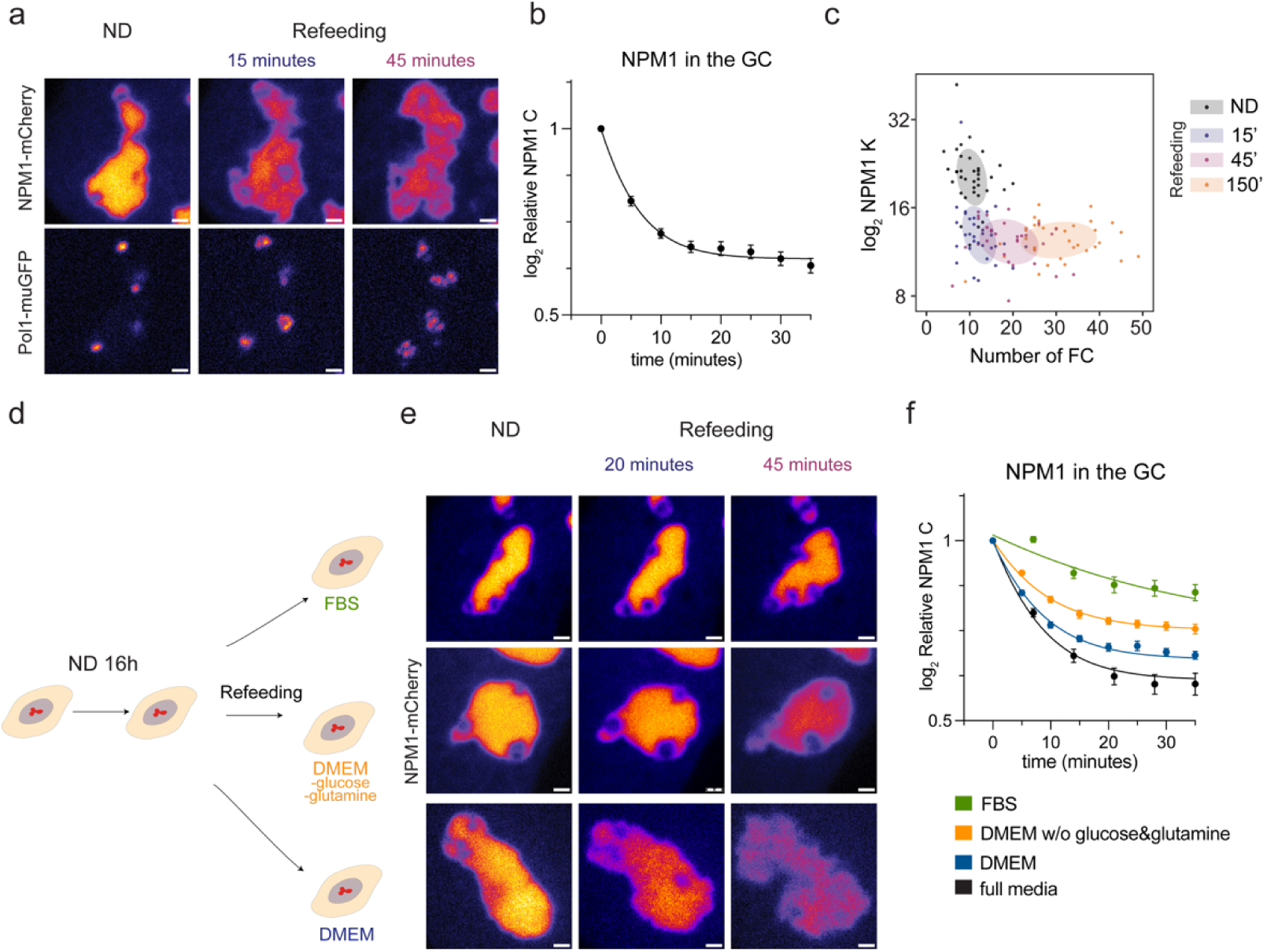
Nutrient re-addition rapidly restores nucleolar properties. (**A**) Dual-edited U2OS cell refed with full media and imaged over time. (**B**) Relative NPM1 levels in the GC over time after media re-addition; normalized to pre-refeeding values; the same cells were tracked throughout the experiment (N = 18 cells); data are shown as mean ± S.E.M.; line indicates one-phase exponential fit (**C**) Scatter plot of NPM1 partitioning versus the number of FCs per cell; N ≥ 29 cells per time point; shaded regions correspond to 68% confidence region. (**D**) Schematic of experiment design for E-F. (**E**) U2OS cells with NPM1-mCherry refed either with ND media with addition of 10% FBS, DMEM, or DMEM without glucose and glutamine and imaged over time. (**F**) Relative NPM1 levels in the GC over time (as in B); N ≥ 15 cells per condition; lines indicate one-phase exponential fit. All images are enlarged views of the nucleolus, displayed in fire LUT, with 1 µm scale bars; display ranges were fixed across time points using the maximum intensity of the ND condition.

To better understand the mechanism of restart, we considered whether these changes result from the resumption of rRNA transcription. We tracked the increase in FC number or nucleolar area, both strong reporters of rRNA transcription activity, and found no significant (p = 0.25) change in the number of FC during the first 15 min of restart. Full recovery of FC number required more than two hours, in contrast to the near-complete recovery of GC NPM1 levels within minutes (**Fig. 4C; Fig. S4C**). Strikingly, these data indicate that rRNA transcription can be functionally uncoupled from GC properties associated with rRNA folding and maturation^3,20^ and more broadly that cells can rapidly remodel nucleolar properties in response to nutrient fluctuations. Thus, these data present nucleolar hibernation as a reversible physiological adaptation to nutrient stress.

Because nucleolar hibernation is tightly coupled to nutrient availability, we next asked whether recovery depends on restoration of cellular energy metabolism. We performed refeeding experiments with partial re-supplementation of media components (**Fig. 4D**). Restoring FBS alone to the usual concentration of 10% led to a small degree of recovery with NPM1 levels decreasing to 0.75 relative to the pre-refeeding baseline after 30 min, while re-supplementing cells with full media triggered a stronger recovery to 0.58 (**Fig. 4E-F**). In cell culture media, the major substrates for cellular energy are glucose and glutamine supporting glycolysis and oxidative phosphorylation; we therefore examined nucleolar recovery without these key energy sources. Refeeding with DMEM not containing glucose and glutamine attenuated recovery with levels dropping to 0.71 versus 0.62 for regular DMEM. Together, these results suggest that resumption of normal nucleolar organization and activity relies on broader metabolic restoration rather than a single nutrient input (**Fig. 4E-F**).

### Nucleolar hibernation is driven by cellular ATP levels

The dependence of nucleolar recovery on nutrient supply suggests that nucleolar hibernation is closely tied to cellular energy status. Because rRNA processing relies on multiple ATP-dependent remodeling and maturation steps^35^, we next asked whether cellular ATP levels regulate nucleolar hibernation. Treatment of cells with Oligomycin A, an inhibitor of mitochondrial ATP synthase, phenocopied nucleolar hibernation in RPE1 cells, as assessed by nucleolar morphology and NPM1 levels in the GC (**Fig. 5A-B**). Additionally, analysis of rRNA intermediates by qPCR revealed processing profiles similar to those observed under ND conditions (**Fig. 5C**). Likewise, ATP depletion reduced the Da number as reflected by the distance at which 50% of MS2-tagged transcripts remained within the nucleolus (0.61 ± 0.01 μm) and final exponential decay slope (2.8 ± 0.1 μm^-1^) in line with a decrease in diffusivity of MS2-tagged rRNA both within and between nucleoli, closely recapitulating the transport defects observed during nucleolar hibernation (**Fig. 5D**). Together, these findings indicate that decreases in ATP levels are sufficient to drive nucleolar hibernation.

**Figure 5.**
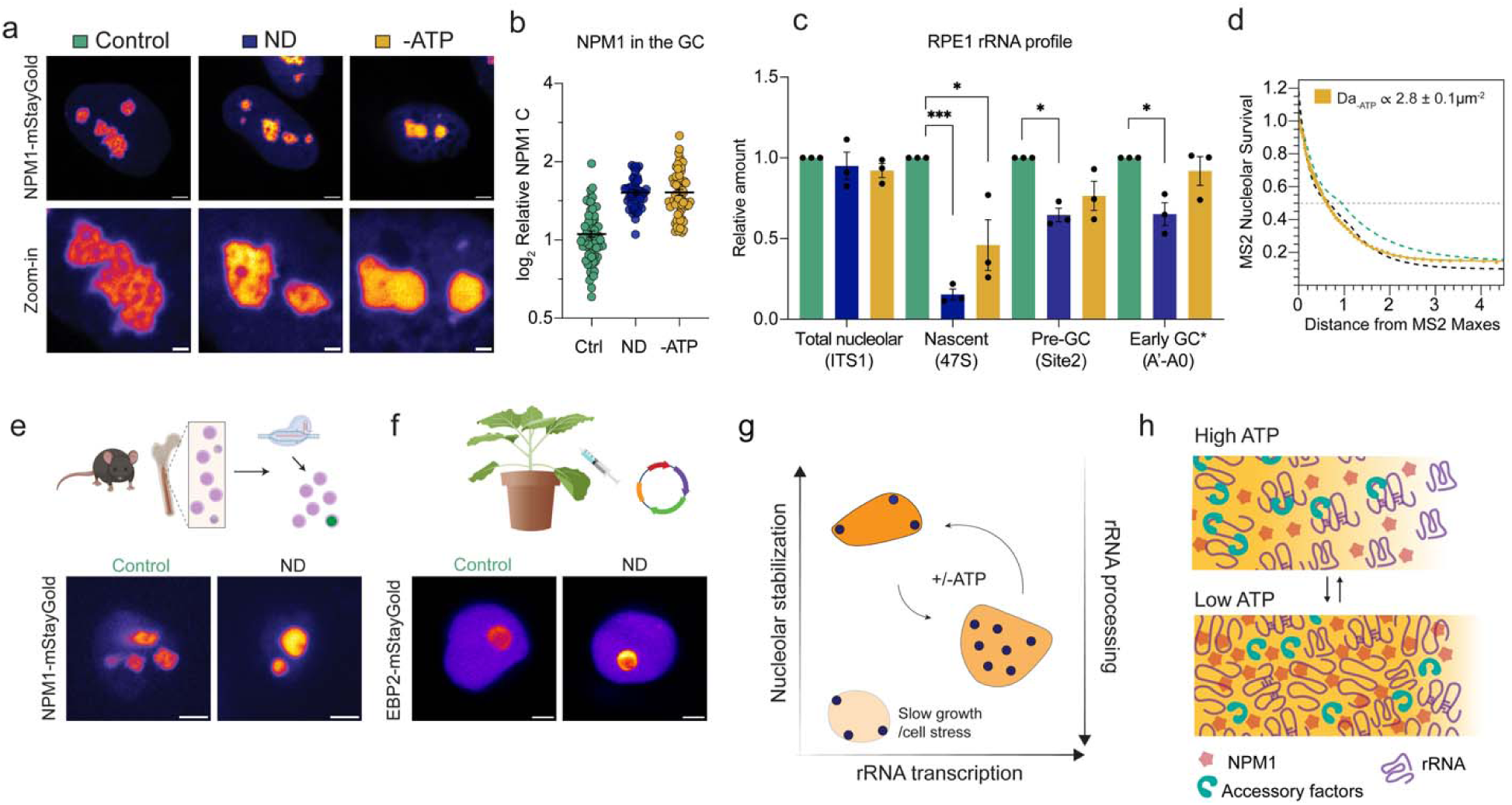
Cellular energy levels tune a conserved hibernating nucleolar state. (A) Live-cell imaging of RPE1 HTERT cells with endogenously edited NPM1-mStayGold; control, nutrient deprivation, and Oligomycin A treatment; scale bar 3 µm (top), 1 µm (bottom). (B) Relative levels of NPM1 in the GC in cells from (A), normalized to the median of the control condition, N ≥ 49 cells per condition; data are shown as mean ± S.E.M. (C) qPCR measurement of relative levels of rRNA intermediates, normalized to control (mean fold change ±S.E.M.; paired t test, n = 3; *p<0.05). (D) MCP-Halo labeled rRNA signal decay within the nucleolus as a function of distance from each position of maximum intensity, averaged and fit; N ≥ 20 cells for Oligomycin A treated cells. (E) CRISPR/Cas9 knock-in of NPM1-mStayGold in mouse hematopoietic stem cells, images for control and ND cells, scale bar 3 µm. (F) Overexpression of EBP2-mStayGold in N. benthamiana leaf epidermal cells, images for control and ND cells, see methods for nutrient-deprived conditions; scale bar 3 µm. (G) Proposed model of nucleolar states. Serum deprivation and nutrient deprivation both suppress rRNA transcription but have opposite effects on GC properties. ATP-sensitive nucleolar hibernation is characterized by reduced rRNA processing and enhanced GC stabilization despite low transcriptional activity. (H) Working model summarizing the proposed mechanism of nucleolar hibernation. Nutrient and ATP limitation slow rRNA processing, resulting in retention of rRNA intermediates, altered transport dynamics, and an overall denser GC.

Energy limitation is a universal challenge faced by all organisms and, across eukaryotes, the role of the nucleolus in ribosome biogenesis is conserved. We therefore asked whether adaptive tuning of nucleolar phase separation extends beyond human cell lines to other eukaryotic cells. First, we edited endogenous NPM1 with mStayGold in primary mouse hematopoietic stem cells and subjected them to nutrient deprivation. In response to ND, NPM1 levels in the GC increased, consistent with human cell lines data (**Fig. 5E, Fig. S4C**). Next, we assessed nucleolar response to ATP depletion in *N. benthamiana* leaves by transiently overexpressing the accessory factor EBP2 fused with mStayGold and subjecting plant leaves to Oligomycin A treatment. Similarly, this energy deprivation resulted in an increase of EBP2 levels in the nucleolus (**Fig. 5F, Fig. S4D**). In both cases, morphological changes mirrored nucleolar hibernation phenotypes suggesting similar cellular responses. Together, our findings reveal nucleolar hibernation as a conserved adaptive state that couples cellular energy availability to ribosome biogenesis.

## Discussion

Cellular metabolic activity dictates ribosome demand, with increased ribosome production reflected by elevated RNA polymerase I activity and enlarged nucleoli^36,37^. Diverse stress responses, including response to starvation, are often interpreted within this framework: stress decreases ribosome biogenesis which results in decreased nucleolar size. The emergence of phase separation as a biophysical concept has added depth to this view by suggesting that nucleolar morphology and size are not simply readouts of transcriptional status but could also reflect composition-dependent stability, material properties, and nanoscale microenvironments that support ribosome biogenesis^15–18^. Indeed, perturbations such as drug treatments, protein overexpression, genetic disruption, and optogenetic gelation have shown that altered nucleolar biophysical properties impairs rRNA processing^4,5,19,23,38^. However, these approaches force or disrupt nucleolar organization and may not capture the coordinated responses of cells to physiological stimuli. Here, we asked whether nutrient stress responses tune more than nucleolar size. We find that nucleoli are not fixed entities, but can be dynamically remodeled in composition, transport properties, and material state.

Quantitative live-cell imaging revealed that nutrient deprivation (ND) and serum deprivation (SD) both decrease nucleolar size and FC activity, yet have opposite effects on GC stabilization. SD effects follow the intuitive slow growth trajectory, in which reduced rRNA transcription is accompanied by smaller nucleoli that expels NPM1 into the nucleoplasm decreasing its partitioning. ND instead induces a hibernating nucleolar state in which reduced rRNA transcription and smaller nucleoli are coupled to increased NPM1 GC levels and partitioning, retention of rRNA intermediates, restricted rRNA transport, reduced diffusivity, and a denser NPM1-proximal microenvironment. Thus, reduced ribosome biogenesis is not a single nucleolar state, and transcriptional output and GC stabilization are not obligatorily coupled. Rather, nucleolar regulation separates into at least two axes (**Fig 5H**). This distinction suggests more complex cellular decision-making, where different stresses may require different ways of slowing ribosome biogenesis. In ND, hibernation may allow cells to slow assembly while sequestering immature ribosomal intermediates within a stabilized GC, preventing premature release when maturation kinetics are delayed (**Fig. 5G**). In this view, cells tune not only how much ribosome biogenesis occurs, but also how intermediates are physically handled.

Our findings extend the emerging model of the nucleolus as an assembly-line condensate whose local microenvironments are coupled to rRNA maturation^15,20–22,24,25^. In active nucleoli, progressive rRNA processing is thought to remodel the GC meshwork, loosening local interaction environments and facilitating the transport and release of maturing ribosomal particles. During hibernation, nutrient and ATP limitation appear to shift this balance by slowing ATP-dependent remodeling steps, retaining earlier rRNA intermediates increasing their valence with GC scaffolding proteins, and stabilizing a denser NPM1-rich microenvironment. The close match between slowed rRNA-processing kinetics, reduced rRNA diffusivity, increased NPM1 partitioning, and decreased local mesh size suggests that nucleolar material properties are coordinated with the biochemical state of ribosome biogenesis. Rapid recovery after nutrient re-addition further argues that hibernation is not failed ribosome biogenesis, but a regulated pause state that poises the nucleolus for restart, consistent with earlier work showing that rRNA intermediates can re-enter maturation after cell refeeding^39^.

The observation of hibernation-like remodeling across divergent systems further suggests a conserved response, although its strength and thresholds likely vary across cell types and physiological contexts. More broadly, our work suggests that condensates are not fixed compartments whose organization is passively dictated by biochemical output. Rather, condensates may function as adaptive materials whose properties are tuned in cellular contexts where energy status and biosynthetic demand are actively remodeled, including during development, differentiation, stress, and recovery.

## Acknowledgements

We thank members of the Riback laboratory for discussions and feedback on the manuscript. We thank Nolwenn Briand for valuable suggestions on the project. This work was supported by funding from the NIH, R35GM162528 (J.A.R.), CA183252 (M.A.G), and CA265748 (M.A.G.). J.A.R. is a CPRIT Scholar in Cancer research with funding from the Cancer Prevention and Research Institute of Texas (CPRIT) New Investigator Grant RR210040 and was supported by the Ted Nash Long Life Foundation, the Leukemia Research Foundation, the Searle Scholars Program, and the Welsh Foundation (Q-2254-20250403). This project was also supported by the Cytometry and Cell Sorting Core at Baylor College of Medicine with funding from the CPRIT Core Facility Support Award (CPRIT-RP180672) and the NIH (P30 CA125123 and S10 RR024574). E.P. was supported by a Marie Skłodowska-Curie COFUND fellowship (Grant Agreement 101126636) during the preparation of this manuscript. E.B. and S.T.H. were supported by a training fellowship from the Houston Area Molecular Biophysics Training Program (NIH T32 GM150582). Y.F. was supported by the George E. Hewitt Foundation for Medical Research Fellowship. H.M. and B.M. were both supported by the Wellcome Trust Investigator award 223049/Z/21/Z to BM. E.K. was supported by the American Cancer Society award (PF-25-1513901-01-PFCBI).

## Author contributions

Conceptualization, E.P. and J.A.R.; Investigation, E.P., E.K., Y.F., E.B., N.C., S.T.H., Z.I., C.D.; Methodology, E.P., H.M., B.M., J.A.R.; Data curation, E.P., J.A.R.; Formal analysis, E.P., E.K., Y.F., J.A.R.; Writing - original draft, E.P., and J.A.R.; Writing - review & editing: all authors; Supervision, L.S., B.M., M.A.G., J.A.R.

## Materials and methods

### Cell culture and nutrient deprivation

All cell lines were cultured in a humidified incubator at 37°C and 5% CO2, in growth media supplemented with 1% penicillin-streptomycin and 10% FBS (Avantor, 89510-186). U2OS cells were cultured in McCoy’s 5A medium (Thermofisher, 16600082); RPE1 HTERT cells were cultured in DMEM/F12 (Thermofisher, 16600082). We routinely tested all cells for Mycoplasma every two weeks using a PCR kit. Forty-eight hours before imaging, cells were plated into glass-bottom 8-well chambers (Ibidi, 80827) coated with fibronectin (Advanced Biomatrix, 103701-196) at a seeding density of 20 000/cm2. For serum deprivation, cells were placed in media with the desired FBS concentration and treated for 48 hours. For nutrient deprivation experiments, we diluted the medium with PBS + 1% BSA to the desired nutrient concentration and treated cells for 16 hours. For refeeding experiments, we replaced nutrient-deprivation media with complete media.

### Generation of CRISPR/Cas9 knock-in lines

We selected guide RNA sequences from the OpenCell database (NPM1, POLR1A) or designed the guide sequence using the Benchling CRISPR tool (Npm1 for mouse); see Supplementary Table 1 for sequence. HDR template sequences were designed using the Benchling HR template tool. Ready-to-use gRNAs were ordered from Synthego with standard modifications. HDR templates were amplified from plasmids containing selected fluorophores using Roche KAPA 2x HiFi HotStart ReadyMix (Roche, 07958935001) in two sequential amplification rounds to extend the homology arms, primers can be found in Supplementary Table 1. HDRs were purified using the AMP Pure bead cleanup protocol to select longer fragments (BD, A63881). For editing, 250,000 cells per reaction were mixed with RNP complexes and HDR sequences - 2 ug Cas9 (IDT, 1081060), 2 ug gRNA, 2 ug HDR fragments - then electroporated using the Lonza Nucleofector SE kit with the following programs: CM-104 for U2OS, EA-104 for RPE1 HTERT, EO-100 for mouse hematopoietic stem cells (see section below). After electroporation, cells were grown in 48-well plates, expanded, and sorted in an isotonic buffer containing 5% FBS using a Sony MA900 sorter. We used at least 3000 cells to establish a stable line, which we further validated by imaging and genotyping PCR.

### RPE1 15/3 28S MS2 cell line generation

RPE1 HTERT cells with an extra copy of chromosome 15, containing MS2 hairpin on 28S rRNA genes were obtained from McStay lab (University of Galway, Ireland). Then, stable expression of MCP-Halo was achieved by lentiviral transduction (lentivirus was a gift from Goodell lab), and NPM1 was tagged with mStayGold fluorophore following the knock-in protocol described above. Prior to imaging, cells were incubated with JaneliaFluor 549 Halo ligand (Promega, HT1020) for 1 hour, then rinsed with pre-conditioned media and imaged. For the analysis of MS2-tagged rRNA gradient, cells having MS2-containing nucleoli made of more than one rDNA chromosome were selected.

### EU pulse-chase

5-EU labeling experiments were performed as described previously^15^. Specifically, cells were grown in an 8-well chamber, treated with nutrient deprivation media for 14 hours, then 1mM 5-EU was added to the cells for 30 minutes and chasing was performed in conditioned nutrient deprivation media from similar cells to avoid changing nutrient concentrations. All the conditions for control and nutrient-deprived cells were processed in parallel so that all samples were fixed together and click reaction (ThermoFisher, C10330) was performed immediately after, followed by imaging.

### Immunofluorescence

Cells were grown in 8-well slide chambers, as for other assays. On the day of fixation, cells were washed 3 times with PBS, fixed with 4% ice-cold paraformaldehyde for 10 min, washed 3 times with PBS, and stored at 4 °C for up to 2 days before staining. For staining, cells were permeabilized in PBS containing 0.1% Triton X-100 (RPI, 111036), 0.01% Tween-20 (Thermo, J20605-AP), and 2% BSA (Roche, 10735078001). Samples were then sequentially incubated with NPM1 antibody (Santa Cruz, sc-32256) and the Alexa Fluor 488 secondary antibody (Jackson Immunoresearch, 711-545-152) for 1 h at room temperature. Slides were kept in PBS at 4 °C after staining and imaged within 2 days.

### NPM1 Local Size Exclusion

For LSE measurements (Figure 3), U2OS NPM1-mCherry were transfected using Lipofectamine 3000 (Sigma) with LSE plasmids encoding NPM1-mStayGold or NPM1-mStayGold with an addition of an amino-acid chain of 200 or 400 AA length. Plasmid sequences can be found in SuppTableX. After 48 hours, cells were subjected to nutrient deprivation and imaged overnight. For details on the LSE method, see ^22^. Specifically, cells were seeded at 20-25,000 per cm^2^ in the 8-well chambers. The next day cells were transfected with 0.25-0.5µg plasmids, 1 µL Lipofectamine P3000 reagent, 0.75 µL Lipofectomine (Thermofisher L3000008), and 25 µL Optimem 1X (Gibco 31985-070) per well. Following 5-hour incubation at 37 °C, transfection media was replaced with complete growth media. After 48 hours, cells were subjected to nutrient deprivation and imaged overnight.

### RNA isolation and qRT-PCR

For RNA isolation cells were seeded into 6-well plates at 20 000/cm^2^ seeding density. RNA was harvested and extracted using Qiagen RNeasy mini kit (Qiagen, 74106). Samples were treated with RNAse-free DNAse on the columns. Concentration was measured using NanoDrop and 1ug of total RNA was used to synthesize cDNA, using Applied Biosystems High-Capacity cDNA Reverse Transcription Kit (4368814). qRT-PCR was performed on a Biorad CFX96 PCR machine using Luna® Universal qPCR Master Mix (NEB, M3003), each reaction was a duplicate. 18S rRNA was used as loading control. All qRT-PCR primer sequences can be found in Supplementary Table 1. Primers were designed using Benchling tool for qRT-PCR primers, sites of pre-rRNA cleavages were validated from^25^. Note that the early-GC primer binds to the nascent rRNA, which is reduced during ND; therefore, the measured difference is underestimated.

### Microscopy

All images were taken on a microscope setup containing a Nikon’s Ti2E, a VisiTech instant SIM microscope (VT-iSIM), a Hamamatsu ORCA Quest qCMOS camera, and a CFI60 Plan Apochromat Lambda D 100X oil immersion objective lens. Cells were maintained in a 5% CO2 at 37°C chamber and imaged with immersion Oil Type B 37°C (Nikon 77005). Imaging of green (muGFP, mStayGold), red (mCherry, JF549), and far-red (Alexa647) was done sequentially via excitation with 488 nm, 561 nm, or 642 nm lasers, respectively. No detectable bleed-through was observed. Power meter measurements were taken at the focal plane for both channels before imaging to verify the proper alignment and performance (e.g., sufficient warming up). Dye standards were also used approximately once a month to determine the no-emission background and vignetting needed for flat field correction for all lasers at all power settings. Additionally, this procedure determined the digital level (DL) offset and the conversion between DL and photoelectrons (via fitting the low-light photon counts histogram) for all camera settings. Linearity of the camera was verified manually as needed. Background field and flat-field corrections, along with the conversion table to relative concentration units (RCU), were applied to the images for quantitative analysis.

### Image analysis

#### ROI selection

Images were analyzed using a custom plugin in Micro-Manager and ImageJ/Fiji. Nuclei were first segmented using the NPM1 channel. Then, to measure NPM1 concentration in the nucleolus, a 4×4-pixel ROI of the maximum intensity was determined using an automated find-maximum procedure. Rectangular ROIs for the nucleoplasm were manually selected. To segment nucleoli for Figure 2 and Figure 3, the NPM1 channel was Gaussian-blurred with a radius of 2 pixels, thresholded using the MaxEntropy method, and subjected to one erosion step. The resulting masks were visually inspected before analysis. FC foci were detected based on Pol 1-muGFP signal and counted, and their centers were used as reference points for radial distribution function analysis within the segmented nucleoli in Figure 2. For MS2 signal survival analysis, RDF centers were instead defined as a 4×4-pixel ROI around the maximum MS2 signal, with nucleolar boundaries determined independently from the thresholded NPM1 channel. RDF analysis was performed as in^15,40^. All automated selections were manually validated and adjusted when necessary.

### 5-EU RDF and transport analysis

Radial distribution function (RDF) analysis was performed as previously described^15,40^ using Pol 1-muGFP-labeled FC centers as radial origins. RDF profiles were fit in Mathematica using the first 888 radial points and a weighted 25th-degree Bernstein polynomial comprising 26 basis functions, with radial position transformed as (x/x^max^)^1.8^, where x^max^ was the final radial position included in the fit. Confidence bands for the fitted curves were calculated from the fit covariance matrix.

Peak positions were determined from the fitted RDF curves. Uncertainty in peak position was estimated by drawing 10,000 parameter sets from the multivariate normal distribution defined by the fitted parameters and covariance matrix and calculating the standard deviation of the resulting peak positions. For comparison between conditions, radial distances were normalized by the corresponding NPM1 peak position. Rescaled RDFs were displayed to 5/3 the relative displacement of NPM1, and RDF intensity was min-max normalized using the extrema of the corresponding fitted curve. Displayed RDF data were logarithmically binned at 100 bins per radial decade.

For quantification of rRNA progression through the GC, normalized 5-EU peak displacement at 0, 30, and 60 min was fit by weighted linear regression using inverse squared peak-position uncertainties as weights. The 135-min time point was excluded from this fit. GC transit time was calculated as the inverse of the fitted displacement rate, with uncertainty calculated using propagation of slope uncertainty used for the reported transit-time error.

Apparent Péclet number was determined from the 135-min 5-EU RDF as previously described^15^. For this analysis, RDF profiles were fit over 5/3 of the NPM1 peak position using the functional form described previously^15^ following fit, data was splined and logarithmically binned as described above.

### MS2

MS2 signal survival was measured as a function of radial distance from the position of maximum MCP-Halo intensity within each MS2-containing nucleolus. Cells were chosen based on the appearance of a nucleolus with non-uniform MS2 signal indicative of containing both the MS2-containing chromosome along with additional rDNA-containing chromosomes. Profiles were pooled across cells and binned for visualization as previously described for RDF analysis ^40^, while nonlinear fitting was performed on the unbinned data over distances of 0–4 µm. Fits were done in Mathematica using NonlinearModelFit to a three-region piecewise analytical solution of the steady-state reaction-diffusion equation ( D (d^2^c/dx^2^) - kc = 0), with fitted transition positions (x_1 and x_2) and reaction-diffusion parameters (k_0, k_1, and k_2). An initial constrained fit was used to determine parameter starting values, followed by nonlinear least-squares fitting using the Levenberg–Marquardt method. The long-distance parameter (k_2) corresponds to k/D = Da/L^2^ and is reported in µm^-2^; parameter uncertainties are reported as standard errors from the nonlinear fit. The distance corresponding to 50% MS2 survival was determined by solving the fitted function for the position at which survival equaled 0.5. Its uncertainty was estimated from the corresponding intersections of the upper and lower mean prediction bands with 0.5 and reported as half the distance between these bounds.

### Generation of NPM1 knock-in mouse hematopoietic cells

Mice were housed in Association for Assessment and Accreditation of Laboratory Animal Care-certified facilities at Baylor College of Medicine and approved by the Institutional Animal Care and Use Committee.

C57BL/6J mice (8-12 weeks old) were initially purchased from the Jackson Laboratory and bred in the facility for the experiments. Hematopoietic stem and progenitor cells (HSPCs) were isolated using ckit+ magnetic beads and recovered for 48 hours in PVA-based cell culture media supplemented with recombinant IL3 and SCF proteins^41^ before gene editing (as described above). mStayGold+ cells were sorted and cultured for up to 7 days in PVA-based cell culture media. Before imaging cells were stained with CD117(APC, 17-1171-83, eBioscience), Ly-6A/E(APC, 17-5981-83, eBioscience) and Mac1 (Pacific Blue, 48-0112-82, eBioscience), Gr1 (Pacific Blue, 48-5931-82, eBioscience), Ter119 (Pacific Blue, 48-5921-82, eBioscience) conjugated antibodies to discriminate immature (CD117 and Ly-6A/E positive) and mature myeloid (Mac1, Gr1, and Ter119 positive) cells. Cells were then subjected to nutrient deprivation (ND) for 2 hours and imaged as described above.

### *Nicotiana benthamiana* EBP2-mStayGold overexpression and nutrient deprivation

*Nicotiana benthamiana* plants were grown under constant light at 22°C. The full-length EBP2 cDNA (AT3G22660) from *Arabidopsis thaliana* fused to mStayGold was cloned into the pTwist ENTR vector by Twist Bioscience and transferred into the pEarleyGate 100 destination vector containing a 35S promoter using LR Clonase II (Invitrogen, 11791020). The resulting construct was introduced into *Agrobacterium tumefaciens* GV3101 by electroporation. Transformed bacteria were selected on LB agar containing rifampicin (15 µg/mL), gentamicin (50 µg/mL), and kanamycin (50 µg/mL), with 1.5% agar, for 2 days at 25°C, followed by overnight growth in liquid LB containing the same antibiotics. Agrobacterium cultures were resuspended in 10 mM MgCl and infiltrated into leaves of 4-week-old plants. Following infiltration, plants were maintained in the dark for 48 h. Before imaging, leaves were infiltrated for 1 h with either 10 µM Oligomycin to deplete ATP or 0.1% ethanol as a vehicle control. A 10 mM oligomycin stock solution (Sigma-Aldrich, O4876) was prepared in ethanol. Abaxial epidermal cells were imaged using a Leica SP8 confocal microscope with a 63× objective, and fluorescence intensities were quantified in Fiji. Nucleoplasmic signal was measured as the mean gray value of a manually placed nucleoplasmic ROI, whereas nucleolar signal was defined as the maximum intensity within the nucleolus after application of a 3 × 3-pixel mean filter.

## Supplementary figures

**Figure S1.**
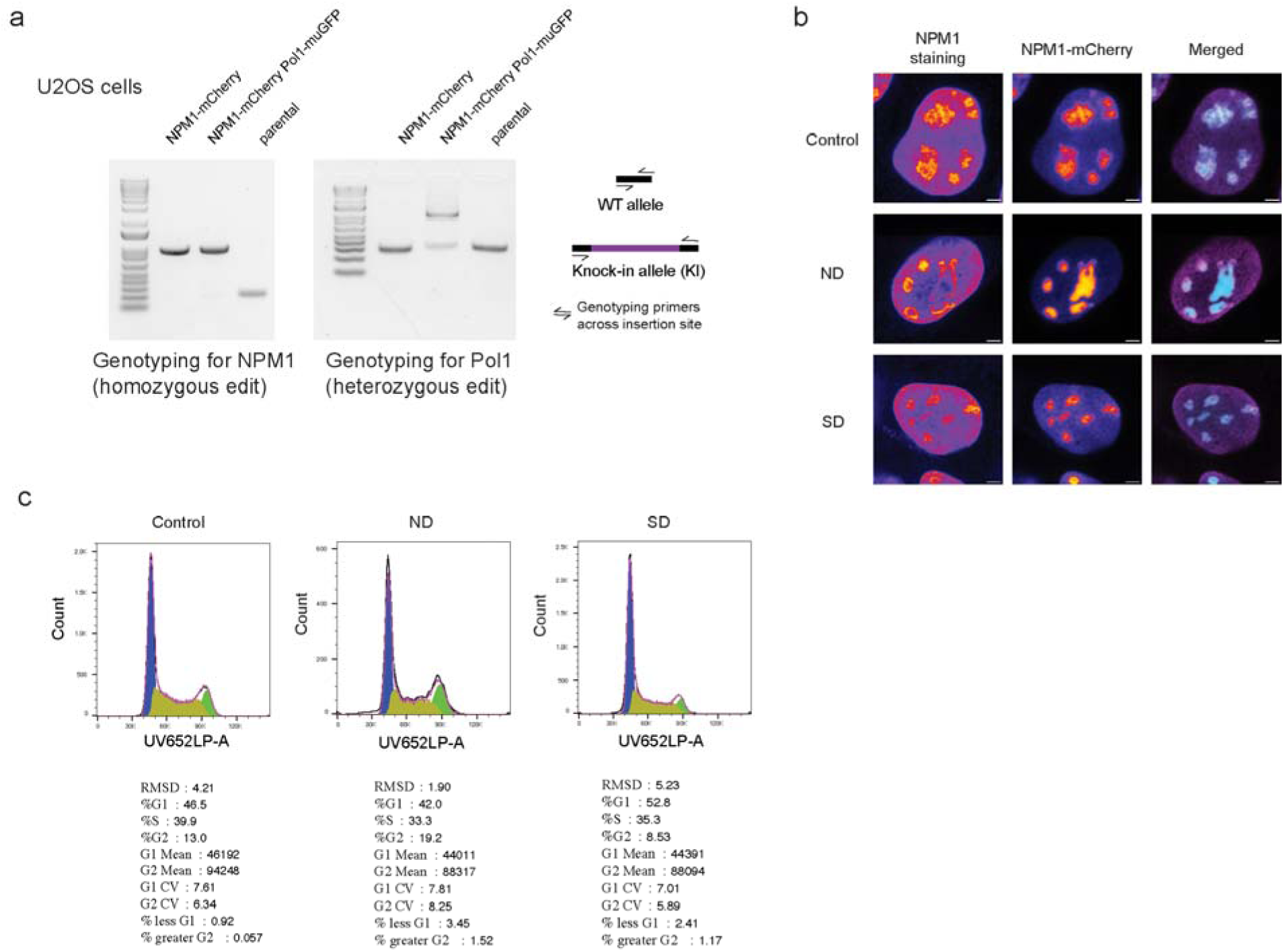
Knock-in line validation and additional controls for nutrient-dependent NPM1 remodeling. (A) Agarose gel electrophoresis for genotyping knock-in lines, PCR across the insertion site. (B) Immunostaining of NPM1 in NPM1-mCherry knock-in U2OS line. Intensities set to its own max values per image. (C) Flow cytometry analysis of cell cycle distribution in U2OS cells, treated with ND and SD and stained with PI.

**Figure S2.**
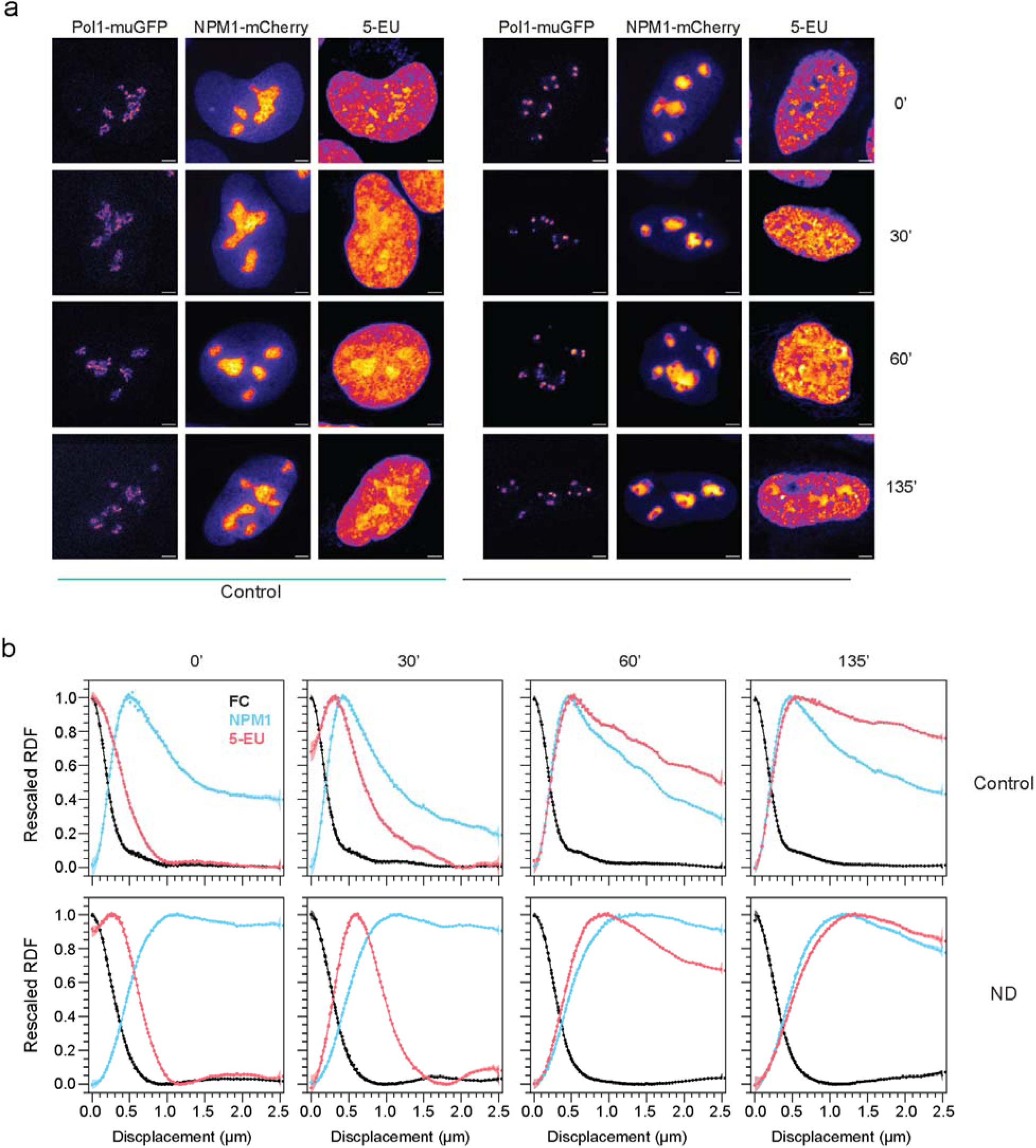
5-EU pulse-chase full nucleus images and RDF plots for individual timepoints with all signals. (A) 5-EU pulse-chase time course in control and ND cells. (B) Correspondingdistribution functions (RDFs); N ≥ 10 cells per time point. radial

**Figure S3.**
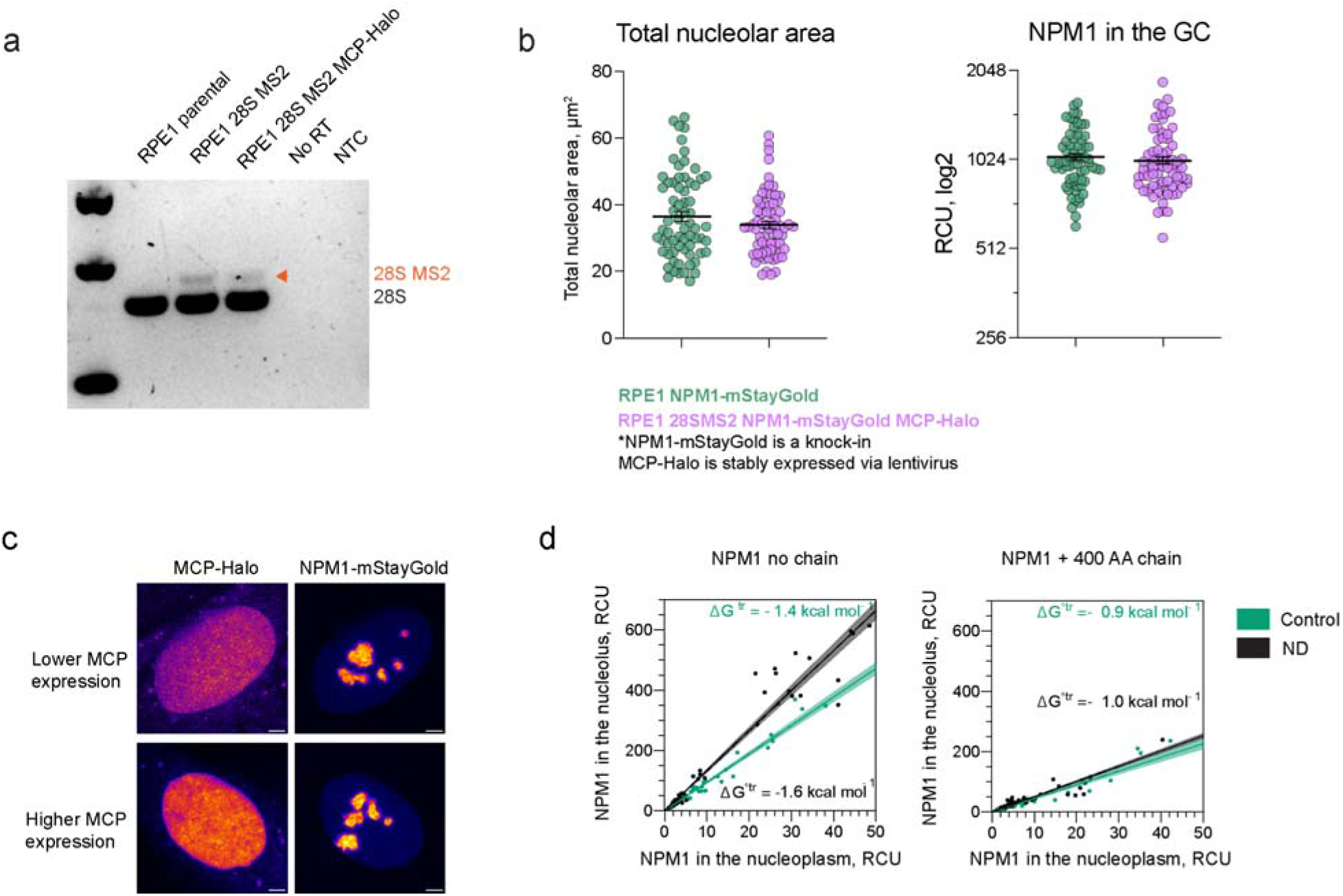
The MS2-containing chromosome does not affect nucleolar parameters in RPE1 cells. (A) Agarose gel electrophoresis of 28S RT-PCR in parental RPE1, RPE1 with the extra chromosome, and with extra chromosome and stable MCP-Halo expression. Arrow marks the 28S-MS2 product, indicating that rRNA with the MS2 hairpin progresses to mature state. (B) Nucleolar parameters of the RPE1 NPM1-mStayGold cells line when compared to the RPE1 28SMS2 NPM1-mStayGold MCP-Halo line. (C) MCP-Halo expression in RPE1 cells not containing the extra chromosome 15, showing MCP does not preferentially localize to the nucleolus. (D) Scatter plot for NPM1 levels n cells expressing NPM1 with no chain and NPM1 fused with 400 AA chain, in control and ND conditions; extrapolation to 0 level expression to determine free energy of transfer (ΔGtr).

**Figure S4.**
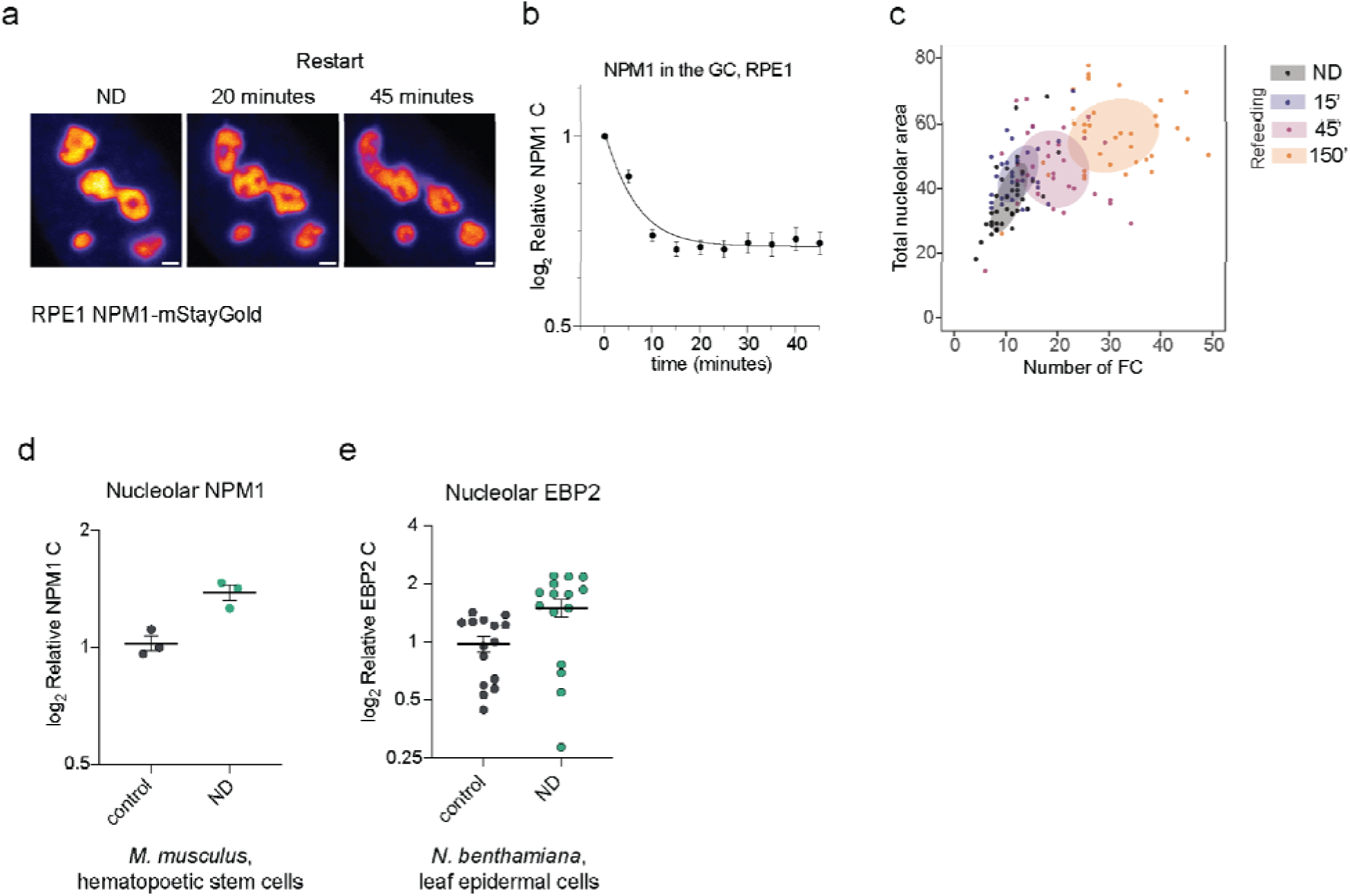
Reversibility of nucleolar hibernation and hibernation-like phenotypes in other systems. (A) Refeeding RPE1 NPM1-mStayGold cells, images of nucleoli for the same cell, ND and refeeding; scale bar 1 µm, display ranges are fixed across time points using the maximum intensity of the ND condition. (B) Relative NPM1 levels in the GC over time after media re-addition; normalized to pre-refeeding values; the same cells were tracked throughout the experiment (N = 10 cells); data are shown as mean ± S.E.M.; line indicates one-phase exponential fit. (C) Scatter plot of total nucleolar area (µm^2^) versus the number of FCs per cell; N ≥ 29 cells per time point; shaded regions correspond to 68% confidence region. (D) Relative NPM1-mStayGold concentration in the nucleolus of mouse HSCs, control and ND; n = 3 cells. (E) Relative EBP2-mStayGold concentration in the nucleoli of *N. bethamiana* abaxial epidermal cells, control and ND, see methods for nutrient-deprived conditions; n = 15 cells.

**Supplementary Table 1.**
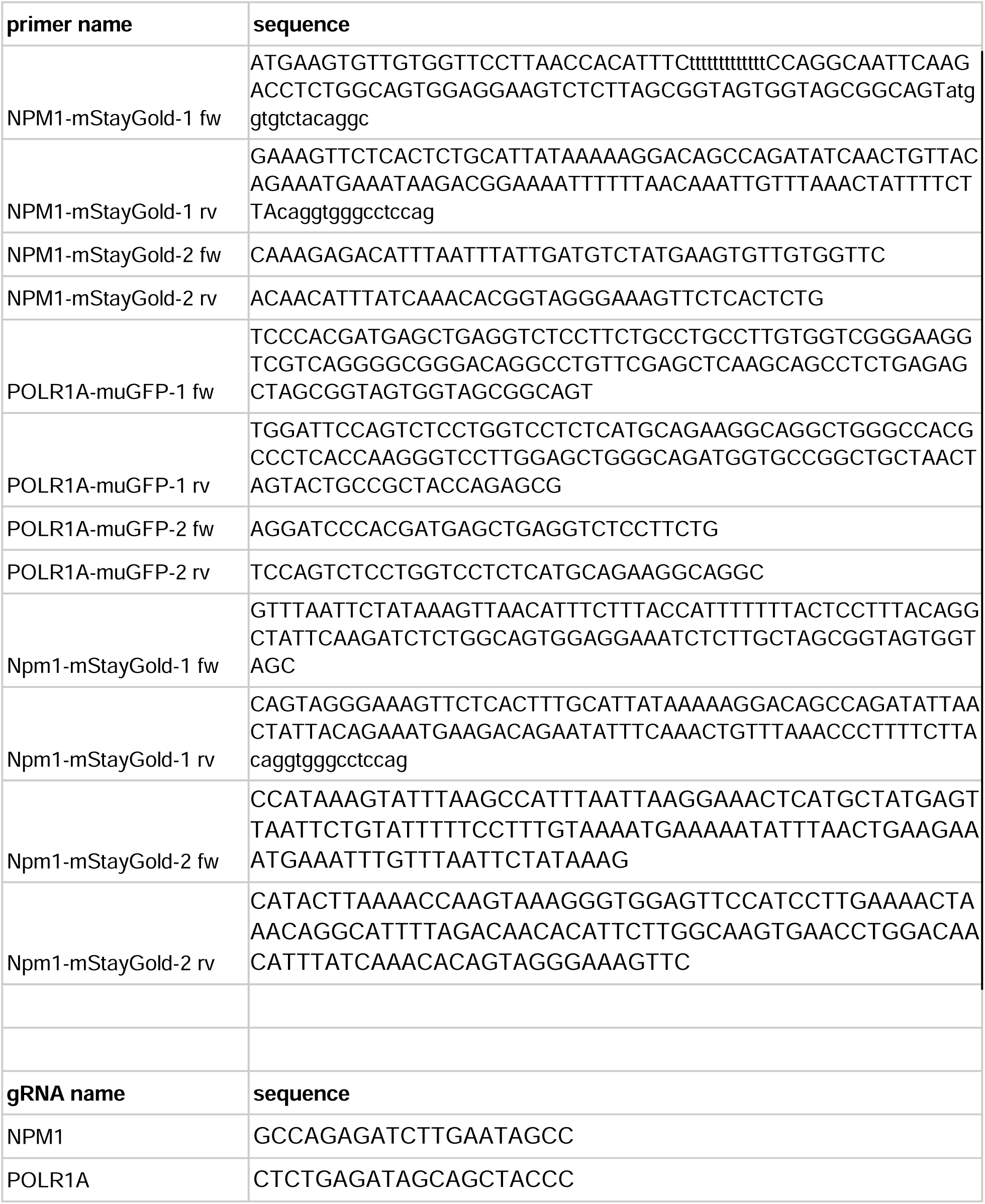
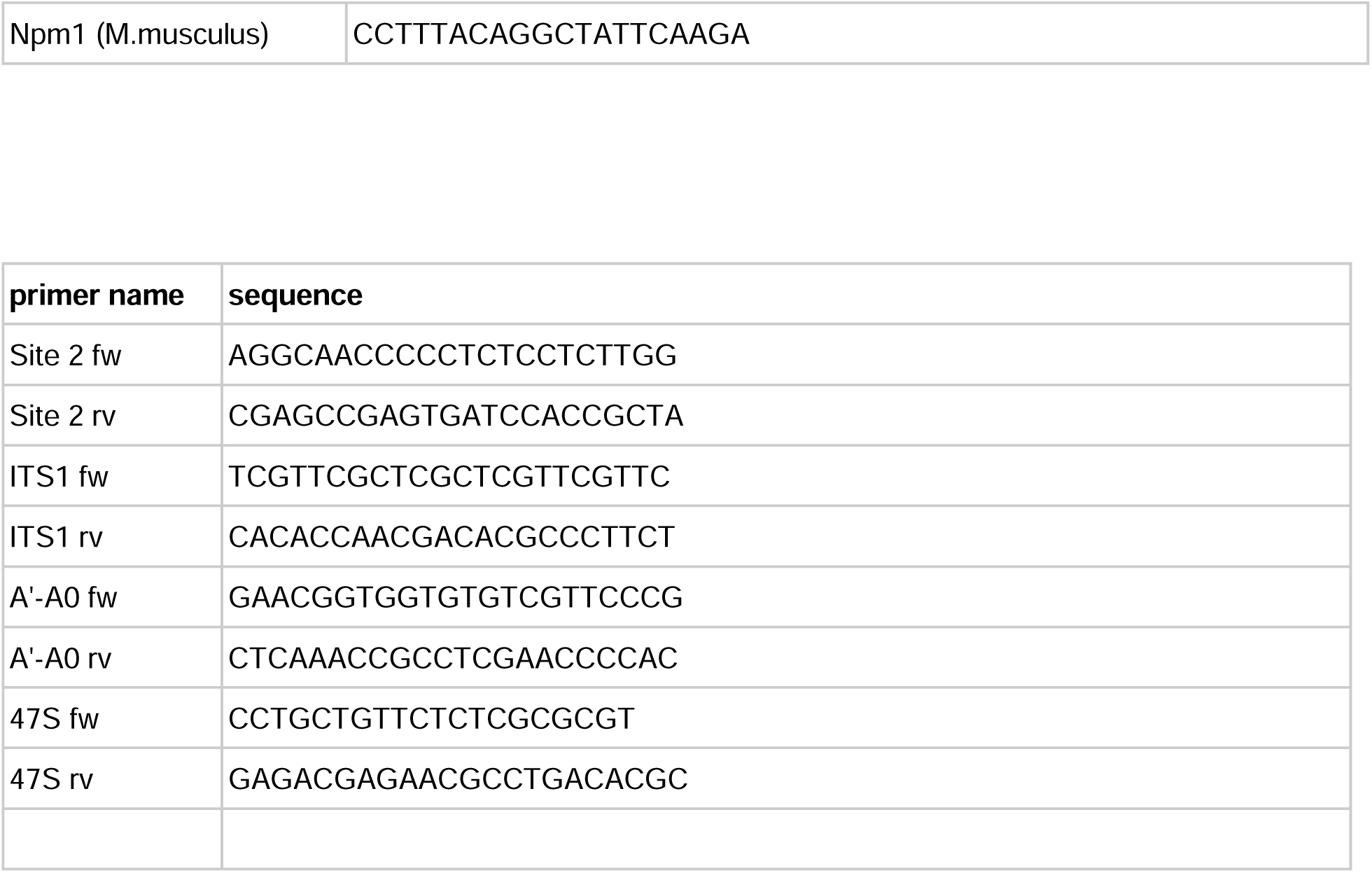

## References

1. Saxton, R. A. & Sabatini, D. M. MTOR signaling in growth, metabolism, and disease. Cell 169, 361–371 (2017).

2. Pakos-Zebrucka, K. et al. The integrated stress response. EMBO Rep. 17, 1374–1395 (2016).

3. Lafontaine, D. L. J., Riback, J. A., Bascetin, R. & Brangwynne, C. P. The nucleolus as a multiphase liquid condensate. Nat. Rev. Mol. Cell Biol. 22, 165–182 (2021).

4. Potapova, T. A. et al. Distinct states of nucleolar stress induced by anticancer drugs. Elife 12, (2023).

5. Zhu, L. et al. Controlling the material properties and rRNA processing function of the nucleolus using light. Proc. Natl. Acad. Sci. U. S. A. 116, 17330–17335 (2019).

6. Boulon, S., Westman, B. J., Hutten, S., Boisvert, F.-M. & Lamond, A. I. The nucleolus under stress. Mol. Cell 40, 216–227 (2010).

7. Burger, K. et al. Chemotherapeutic drugs inhibit ribosome biogenesis at various levels. J. Biol. Chem. 285, 12416–12425 (2010).

8. Spriggs, K. A., Bushell, M. & Willis, A. E. Translational regulation of gene expression during conditions of cell stress. Mol. Cell 40, 228–237 (2010).

9. Pelletier, J., Thomas, G. & Volarević, S. Ribosome biogenesis in cancer: new players and therapeutic avenues. Nat. Rev. Cancer 18, 51–63 (2018).

10. Klinge, S. & Woolford, J. L., Jr. Ribosome assembly coming into focus. Nat. Rev. Mol. Cell Biol. 20, 116–131 (2019).

11. Shore, D. & Albert, B. Ribosome biogenesis and the cellular energy economy. Curr. Biol. 32, R611–R617 (2022).

12. McStay, B. Nucleolar organizer regions: genomic “dark matter” requiring illumination. Genes Dev. 30, 1598–1610 (2016).

13. Yang, K. et al. A redox mechanism underlying nucleolar stress sensing by nucleophosmin. Nat. Commun. 7, 13599 (2016).

14. Yang, K., Yang, J. & Yi, J. Nucleolar Stress: hallmarks, sensing mechanism and diseases. Cell Stress Chaperones 2, 125–140 (2018).

15. Riback, J. A. et al. Viscoelasticity and advective flow of RNA underlies nucleolar form and function. Mol. Cell 83, 3095–3107.e9 (2023).

16. Feric, M. et al. Coexisting Liquid Phases Underlie Nucleolar Subcompartments. Cell 165, 1686–1697 (2016).

17. Cheng, H. H. et al. Micropipette aspiration reveals differential RNA-dependent viscoelasticity of nucleolar subcompartments. Proc. Natl. Acad. Sci. U. S. A. 122, e2407423122 (2025).

18. King, M. R. et al. Macromolecular condensation organizes nucleolar sub-phases to set up a pH gradient. Cell 187, 1889–1906.e24 (2024).

19. Riback, J. A. et al. Composition-dependent thermodynamics of intracellular phase separation. Nature 581, 209–214 (2020).

20. Bose, R. et al. Mapping RNA structure assembly and remodeling in biomolecular condensates. bioRxiv 2026.05. 21.726738 (2026) doi:10.64898/2026.05.21.726738.

21. LaPeruta, A. J., Micic, J. & Woolford, J. L., Jr. Additional principles that govern the release of pre-ribosomes from the nucleolus into the nucleoplasm in yeast. Nucleic Acids Res. 51, 10867–10883 (2023).

22. Dollinger, C. et al. Nanometer condensate organization in live cells derived from partitioning measurements. bioRxivorg 2025.02. 26.640428 (2025) doi:10.1101/2025.02.26.640428.

23. Zhang, Y. et al. Probing condensate microenvironments with a micropeptide killswitch. Nature 1–10 (2025).

24. Pan, Y.-H. et al. Pre-rRNA spatial distribution and functional organization of the nucleolus. Nature 646, 227–235 (2025).

25. Quinodoz, S. A. et al. Mapping and engineering RNA-driven architecture of the multiphase nucleolus. Nature 1–10 (2025).

26. Chen, M. et al. Serum starvation induced cell cycle synchronization facilitates human somatic cells reprogramming. PLoS One 7, e28203 (2012).

27. He, L. et al. Autophagy: The last defense against cellular nutritional stress. Adv. Nutr. 9, 493–504 (2018).

28. de Mesquita, C. B., Kaida, A., Nojima, H. & Miura, M. Reversible nucleolar stress and cell growth arrest triggered by acidic pH. J. Cell. Physiol. 241, e70176 (2026).

29. Ugai, H. et al. Adenoviral protein V promotes a process of viral assembly through nucleophosmin 1. Virology 432, 283–295 (2012).

30. Ochs, R. L. & Smetana, K. Fibrillar center distribution in nucleoli of PHA-stimulated human lymphocytes. Exp. Cell Res. 184, 552–557 (1989).

31. Mangan, H. & McStay, B. Human nucleoli comprise multiple constrained territories, tethered to individual chromosomes. Genes Dev. 35, 483–488 (2021).

32. Deen, W. M. Analysis of Transport Phenomena. (Oxford University Press, New York, NY, 2013).

33. Shen, Y. et al. Biomolecular condensates undergo a generic shear-mediated liquid-to-solid transition. Nat. Nanotechnol. 15, 841–847 (2020).

34. Jawerth, L. et al. Protein condensates as aging Maxwell fluids. Science 370, 1317–1323 (2020).

35. Mitterer, V. & Pertschy, B. RNA folding and functions of RNA helicases in ribosome biogenesis. RNA Biol. 19, 781–810 (2022).

36. Buchwalter, A. & Hetzer, M. W. Nucleolar expansion and elevated protein translation in premature aging. Nat. Commun. 8, 328 (2017).

37. Corman, A., Sirozh, O., Lafarga, V. & Fernandez-Capetillo, O. Targeting the nucleolus as a therapeutic strategy in human disease. Trends Biochem. Sci. 48, 274–287 (2023).

38. Schmidt, H. B. et al. Oxaliplatin disrupts nucleolar function through biophysical disintegration. Cell Rep. 41, 111629 (2022).

39. Pan, M. et al. Glutamine deficiency in solid tumor cells confers resistance to ribosomal RNA synthesis inhibitors. Nat. Commun. 13, 3706 (2022).

40. Datar, G. K. et al. Disparate leukemia mutations converge on nuclear phase-separated condensates. Cell 188, 7118–7136.e21 (2025).

41. Khoo, H. M., Meaker, G. A. & Wilkinson, A. C. Ex vivo expansion and genetic manipulation of mouse hematopoietic stem cells in polyvinyl alcohol-based cultures. J. Vis. Exp. (2023) doi:10.3791/64791.

